# Pulcherrimin Production by Endophytic *Metschnikowia pulcherrima*: Fermentation Performance and Antimicrobial Activity

**DOI:** 10.64898/2026.07.17.739157

**Authors:** Dildora Shodieva, Liliya Abdulmyanova, Toshkhon Gulyamova, Dilorom Ruzieva, Nilufar Kuzieva, Muzaffar Annaev, Shakhnoza Siddikova, Sardor Mahkamov

## Abstract

Endophytic yeasts of the *Metschnikowia pulcherrima* clade are recognized as natural producers of pulcherrimin, an iron-chelating red pigment associated with antimicrobial activity. However, endophytic isolates from arid Central Asian fruit trees remain unexplored, and their pulcherrimin biosynthetic capacity is poorly characterized. In this study, five endophytic *M. pulcherrima* strains were isolated from apricot, persimmon, peach and pomegranate fruits in Uzbekistan and identified using MALDI-TOF MS and ITS sequencing. The strains were cultivated in FeCl₃-supplemented Sabouraud medium, and pulcherrimin was extracted and quantified using a gravimetric workflow with biological triplicates. Pigment yields ranged from 420.6 to 776.4 mg/L, representing some of the highest values reported for non-engineered isolates under flask conditions. Antimicrobial activity was evaluated by agar well diffusion assays against *Staphylococcus aureus*, *Bacillus subtilis*, *Escherichia coli*, *Pseudomonas aeruginosa* and *Candida albicans*, with inhibition zones reaching 18–23 mm depending on the strain. Nano-LC-MS/MS analysis confirmed the presence of key intermediates of the pulcherrimin biosynthetic pathway, including cyclo(Leu–Leu) and pulcherriminic acid, supporting the metabolic origin of the pigment. These findings reveal that endophytic *M. pulcherrima* from Uzbekistan fruit trees and shrubs constitute highly productive natural sources of pulcherrimin and represent promising candidates for future fermentation optimization and development of natural antimicrobial pigments.

**Paragraph on importance:** This study provides the first systematic evidence that endophytic Metschnikowia pulcherrima yeasts isolated from fruit trees in Central Asia possess an exceptional capacity to produce pulcherrimin, a natural antimicrobial pigment. Strains obtained under the environmental conditions of Uzbekistan produced remarkably high levels of pulcherrimin (up to 776 mg/L) under simple, non-genetically modified fermentation conditions, exceeding yields previously reported for yeast and bacterial sources. The iron-chelating mechanism characteristic of pulcherrimin confers broad-spectrum antimicrobial activity against bacteria and yeasts. These findings highlight endophytic yeasts as environmentally safe and sustainable sources of natural antimicrobial pigments and identify Central Asia as a previously unexplored reservoir of high-value microbial resources with significant biotechnological potential.

## Introduction

*Non-Saccharomyces* yeasts have gained increasing attention in biotechnology and sustainable agriculture due to their unique metabolic capabilities and antagonistic properties [1,2]. Among them, *Metschnikowia pulcherrima* stands out for its ability to produce pulcherrimin, a reddish iron-chelating pigment derived from pulcherriminic acid, which exerts strong antimicrobial activity primarily through iron sequestration in the extracellular environment [3,4]. This mechanism enables *M. pulcherrima* to effectively outcompete a broad range of fungal pathogens and bacteria, including major postharvest spoilers such as *Botrytis cinerea, Penicillium expansum, Monilinia spp*., and food-borne pathogens like *Listeria monocytogenes* and *Salmonella enterica* [5,6,7,8]. Pulcherrimin itself exhibits antioxidant, photoprotective, and potential cosmetic applications, broadening its industrial relevance [3,9,10,11,13].

Pulcherrimin biosynthesis is governed by the PUL gene cluster, with PUL1 and PUL2 encoding the core cyclodipeptide oxidase and synthetase, respectively [12,14,15]. Environmental factors, especially iron limitation and the presence of specific surfactants (e.g., Tween-80), dramatically enhance PUL gene expression and pigment yields, reaching up to 300–330 mg/L in optimized fermentation conditions [16]. Recent heterologous expression studies in *Saccharomyces cerevisiae* and metabolic engineering efforts have further confirmed the central role of PUL1/PUL2 and identified rate-limiting steps in the pathway [12,17].

Despite significant progress, most characterized high-pulcherrimin producers originate from vineyard-associated environments in Europe, North America, and East Asia [13,18,19]. Endophytic isolates, particularly from arid and semi-arid regions, remain largely unexplored, even though plants growing under abiotic stress are known to host highly competitive microbial communities with enhanced secondary metabolite production [20,21]. Central Asia, with its extreme continental climate and rich fruit biodiversity, represents a promising yet untapped reservoir of novel *M. pulcherrima* strains.

In this study, we isolated and characterized five endophytic *M. pulcherrima* strains from traditionally cultivated fruit trees and shrubs (apricot, persimmon, peach and pomegranate) in Uzbekistan. Using a combination of morphological, molecular (ITS sequencing), proteomic (nano-LC-MS/MS), and fermentation approaches, we evaluated strain-dependent pulcherrimin production capacity (420.6-776.4mg/L), confirmed pigment identity and associated metabolome, and assessed antimicrobial activity against relevant postharvest and clinical pathogens. This work provides the first comprehensive insight into pulcherrimin-producing endophytic *M. pulcherrima* from Uzbekistan fruit trees, shrubs and highlights their potential as novel sources of natural iron-chelating antimicrobials for sustainable postharvest protection and biotechnological applications.

## 2. Materials and Methods

### 2.1. Isolation Endophytic Yeast Strains

Endophytic yeasts were isolated from fresh fruits of plants using the method described by Abdel-Hafez et al. [23]. Fruits of plants growing in the Yunusobod district of Tashkent city, Republic of Uzbekistan — *Prunus armeniaca* (apricot), *Diospyros kaki Tpip*. (persimmon), *Prunus persica* (peach), and *Punica granatum* (pomegranate)— were collected and the isolates were obtained at the Laboratory of “Biochemistry and Biotechnology of Physiologically Active Compounds” of the Institute of Microbiology, Academy of Sciences of the Republic of Uzbekistan. All plant materials were identified based on their morphological characteristics.

The samples were processed under aseptic conditions. Surface sterilization was carried out using 70% ethanol (3–5 min), followed by treatment with 3% sodium hypochlorite, and subsequently rinsed three times with sterile distilled water. The final rinse water was plated onto agar medium as a sterility control, and no microbial growth was observed, confirming effective surface sterilization.

The sterilized materials were cut into fragments of approximately 5 mm. The plant fragments were then transferred into flasks containing liquid Sabouraud medium supplemented with cefotaxime (200 mg/L) to suppress bacterial growth during the initial isolation stage. After the primary enrichment step, cefotaxime was not used in any subsequent cultivation procedures. The cultures were incubated at 28 °C, 140 rpm for 3–5 days on a rotary shaker.

After visual changes such as turbidity and gas formation were observed, the enriched cultures were streaked onto Sabouraud Dextrose Agar (SDA) plates without antibiotics to obtain pure cultures. Yeast-like colonies were microscopically examined and repeatedly sub-cultured on SDA to ensure purity.

The obtained isolates were preliminarily identified according to the methods of Abdel-Hafez et al. (2015) and Kurtzman et al. (2000). For the separation of the culture liquid (supernatant), the yeast cultures were centrifuged at 6000 rpm for 15 minutes. The supernatant was collected for further analyses. The obtained isolates were preliminarily classified according to the criteria described by Kurtzman et al. [24].

### 2.2. Morphological and Molecular Identification of Yeasts

Initially, identification was performed using the MALDI-TOF method. MALDI-TOF MS (Matrix-Assisted Laser Desorption/Ionization Time-of-Flight Mass Spectrometry) [25] was followed by molecular identification.

Identification was carried out by MALDI-TOF and by DNA amplification and sequencing.

For preliminary conventional classification, the shape, colour, and size of yeast colonies as well as the morphological structure of the cells were examined. For this purpose, 3–5-day-old yeast colonies grown on Petri dishes were used.

Subsequently, 48–72-hour yeast isolates grown as single colonies on Sabouraud medium were used for identification by MALDI-TOF mass spectrometry. MALDI-TOF MS is a widely applied and common method in clinical microbiological laboratories for the identification of pathogenic bacteria, especially microaerophiles, anaerobes, mycobacteria, and fungi.

Identification of *Metschnikowia pulcherrima* ELM-GS-3 by Sequence Analysis of the Internal Transcribed Spacer (ITS) Regions For DNA isolation, the EurX GeneMATRIX Plant & Fungi DNA isolation kit (Poland) was used. After DNA isolation, the quantity and purity of the obtained DNA were assessed using the Thermo Scientific Nanodrop 2000 (USA) spectrophotometer. For the PCR analysis, the universal primers ITS1 and ITS4 were used to amplify the target gene regions for species identification. The primer sequences were as follows: ITS1 5’ TCCGTAGGTGAACCTGCGG 3’ and ITS4 5’ TCCTCCGCTTATTGATATGC 3’. PCR conditions included an initial denaturation at 95°C for 5 minutes, followed by 40 cycles of denaturation at 95°C for 45 seconds, annealing at 57°C for 45 seconds, and extension at 72°C for 60 seconds. A final extension was performed at 72°C for 5 minutes, and the temperature was then lowered to 4°C to complete the PCR. The amplification products obtained with the Kyratec thermocycler were analyzed by electrophoresis on a 1.5% agarose gel prepared with 1x TAE buffer, run at 100 volts for 90 minutes, and visualized under UV light using ethidium bromide staining. A single-step PCR was performed to amplify a region of approximately 700 base pairs, using Solis Biodyne (Estonia) FIREPol® DNA Polymerase Taq polymerase enzyme. The presence of a single band on the agarose gel indicated successful PCR amplification. For PCR product purification, the single-band samples were purified using the MAGBIO “HighPrepTM PCR Clean-up System” (AC-60005) according to the manufacturer’s procedures. Sanger sequencing was carried out at Macrogen’s laboratory in the Netherlands, using the ABI3730XL Sanger sequencing device (Applied Biosystems, Foster City, CA) and the BigDye Terminator v3.1 Cycle Sequencing Kit (Applied Biosystems, Foster City, CA). The reads obtained with ITS1 and ITS4 primers were assembled into a contig to create a consensus sequence using the CAP contig assembly algorithm in BioEdit software.

Some of the identified strains were confirmed using modern genetic methods. This was achieved by sequencing the 18S rRNA, ITS1, and ITS2 gene regions. Initially, yeast samples were grown for 48–72 hours in liquid nutrient medium, and samples were collected for DNA isolation; DNA was obtained through extraction. Polymerase chain reaction (PCR) was employed to obtain partial and complete sequences of the 18S rRNA, ITS1, and ITS2 regions. For this purpose, specific primers targeting the aforementioned genes and the ITS (Internal Transcribed Spacer) regions were selected.

### 2.2 Preparation of Culture Media and Biomass Production

To obtain pulcherrimin pigment, different nutrient media were used as required culture media, and these directly affected the amount of biomass. In particular, pigment production could occur in either nutrient-rich (Sabouraud) or minimal (1% glucose, 0.3% (NH₄)₂SO₄, 0.1% KH₂PO₄, 0.05% MgSO₄ × 7H₂O, 0.05% yeast extract) media [16]. FeCl₃ at 0.05% was additionally added to these media [16]. Yeast strains were cultivated for 5 days at 28°C with shaking at 150 rpm in various volumes for the purpose of pigment production.

### 2.3 Extraction of Pulcherrimin Pigment

Pulcherrimin pigment was extracted from yeast cultures using the method previously described by Kluyver and co-workers[1], later modified by Li and co-workers [19,28]. A 50 mL aliquot of each yeast culture was centrifuged at 4 °C and 5000 × g for 10 min using an Eppendorf 5804R centrifuge (Hamburg, Germany). The pellet containing yeast cells and pigment was treated at 4 °C with methanol (50 mL of 99.8% methanol per 10 g wet yeast biomass). After overnight incubation, yeast cells were centrifuged (4 °C, 5000 × g, 10 min) and subsequently washed twice with distilled water (25 mL). The yeast biomass was resuspended in 2 M NaOH and centrifuged (4 °C, 5000 × g, 10 min). The pH of the supernatant was adjusted to 1.0 with 4 M HCl, and the mixture was incubated at 100 °C for 30 min. The precipitated pigment was collected by centrifugation (4 °C, 8000 × g, 20 min) and washed three times with 25 mL of distilled water.

To obtain pure pulcherrimin, the dissolution–precipitation steps with NaOH and HCl were repeated three times. Finally, the pigment was recovered by centrifugation and dried at 60 °C for 18 h (MA 50.R moisture analyser, Radwag, Poland). The purity of the obtained red pigment was verified by ¹H NMR spectroscopy. The isolated product was dissolved (5% w/v) in deuterated hydroxide solution (2 M NaOH in D₂O) at 25 °C. The liquid-state ¹H NMR spectrum was recorded at 25 °C on a Bruker DRX-500 MHz spectrometer (Billerica, MA, USA).

### 2.4 Antimicrobial Activity

The antimicrobial activity of pulcherrimin pigment isolated from the endophytic yeast *M. pulcherrima* was determined using the agar well diffusion method [7]. The antimicrobial activity of pulcherrimin pigment obtained from the endophytic yeast *M.pulcherrima* was evaluated on meat-peptone agar (GPA) medium by the agar well diffusion method. For the assay, Gram-positive bacteria *Staphylococcus aureus* and *Bacillus subtilis*; Gram-negative bacteria *Pseudomonas aeruginosa* and *Escherichia coli*; and the yeast *Candida albicans* were used. Bacterial strains were grown in 10 mL of meat-peptone broth (GPB) at 37 °C for 18–20 hours in a thermostat. The yeast *Candida albicans* was cultivated in 10 mL of malt extract broth (MEB) at 25 °C for 48 hours. The resulting cultures were diluted 100-fold, and suspensions were then adjusted to the McFarland 0.5 standard (approximately 1 × 10⁷ CFU/mL). Bacterial suspensions (0.1 mL) were spread evenly onto GPA Petri dishes and incubated at 37 °C for 30 minutes. The yeast suspension was similarly spread onto malt extract agar (MEA) Petri dishes and incubated at 25 °C for 30 minutes. After the surface had dried, wells of 10 mm diameter were made. Each well was filled with 100 μL of a 3% (w/v) aqueous suspension of pulcherrimin pigment (dissolved in DMSO at 1 mg/mL). The experiment was performed in triplicate. Cefazolin (1 mg/mL) and clotrimazole (1 mg/mL) were used as positive controls, and DMSO was used as the negative control. Bacterial plates were incubated at 37 °C for 24 hours, and yeast plates were incubated at 25 °C for 7 days. At the end of incubation, the diameters of the inhibition zones (mm) were measured manually, and mean values were calculated. Each experiment was repeated three times.

### 2.5 Identification and characterization of samples using reverse phase Nano-LC-MS/MS

Reverse phase nano-liquid chromatography tandem mass spectrometry (nano-LC-MS/MS) analysis was performed on an Agilent 1200 nano-flow liquid chromatography system. The system was connected to an Agilent 6520B Q-TOF mass spectrometer, with sample fractions separated using a Zorbax SB C18 CHIP column (5 µm, 75 µm x 43 mm). The purpose of the analysis was to identify and characterize analytes in the sample. Samples were dissolved in 50 mM ammonium bicarbonate buffer (pH 8.0). Protein samples were digested with trypsin enzyme (1:50 enzyme:protein ratio) at 37°C for 16 hours. Digested peptides were prepared at 1 µg/µL concentration in 0.1% formic acid and 5% acetonitrile solution. The sample was filtered using a 0.22 µm syringe filter and stored at 4°C before injection. Chromatography conditions Column: Zorbax SB C18 CHIP (5 µm, 75 µm x 43 mm). Mobile phase: A: 0.1% formic acid + 5% acetonitrile; B: acetonitrile + 0.1% formic acid + 10% deionized water. Flow rate: Sample loading: 4.0 µL/min (Agilent 1260 Cap Pump); Elution: 0.6 µL/min (Agilent 1260 Nano Pump). Gradient schedule: B solution concentration was changed linearly as follows: Time (min) B solution (%) 0–3 0 12 60 18 60 20 0 Injection: 2.0 µL sample was injected into the column using Agilent Micro WPS autosampler. Degassing: Mobile phase solutions were degassed using Agilent 1260 µ-degasser. Eluted fractions were analyzed on Agilent 6520B Q-TOF mass spectrometer under the following conditions. Ionization source: Electrospray ionization (ESI), positive ionization mode (ESI+); Drying gas: Nitrogen, flow rate 4.0 L/min, temperature 350°C; Voltage settings: Skimmer cone: 65 V; Fragmentor: 175 V; Capillary voltage: 2000 V (optimized in 1800–2500 V range); Mass range: MS mode: 50–3000 m/z; MS/MS mode: 50–2500 m/z; Fragmentation: Collision energy for ion fragmentation in MS/MS mode was 10–40 eV, optimized based on analyte m/z. Data acquisition: In automatic MS/MS mode, the 4 highest intensity precursor ions were selected, with data collected at 2 spectra/second per ion. System calibration and quality control The mass spectrometer was calibrated before each analysis using Agilent ESI calibration mixture (m/z 118–2722). To check system stability, a quality control (QC) sample (1 ng/µL trypsin-digested BSA) was used after every 10 samples. Caffeine at 100 fmol/µL was added as internal standard. Mass spectrometry data were processed in Agilent MassHunter Qualitative Analysis software (version B.07.00). Peptide searches were conducted against the UniProt database using Mascot and Spectrum Mill search engines. Search parameters: Mass match error: ±20 ppm (MS), ±0.05 Da (MS/MS); Missed cleavages: up to 2; Modifications: cysteine carbamidomethylation (fixed), methionine oxidation (variable).

### 2.6 Statistical Analysis

All experiments were performed in biological triplicates (n = 3). Results are presented as mean ± standard deviation (SD). Comparisons among multiple groups (different strains and/or culture conditions) were analyzed using one-way or two-way analysis of variance (ANOVA) depending on the experimental design. When significant differences were detected (p < 0.05), Tukey’s honestly significant difference (HSD) post-hoc test was applied for pairwise comparisons. Normality and homogeneity of variance were verified prior to ANOVA. Statistical analyses were carried out using Python 3.12 with the statsmodels (v0.14.0) and scipy libraries. Graphs and figures were prepared using Matplotlib and Seaborn

## 3. Results

### 3.1 Isolation and Identification of Endophytic Yeast Strains

To isolate endophytic yeasts, fruits of local plants growing in the territory of the republic were selected (Table 1). From the fruits of *Prunus armeniaca* (apricot), *Diospyros kaki* (persimmon), *Prunus persica* (peach), and *Punica granatum* (pomegranate), a total of five isolates were obtained. These isolates were initially subjected to morphological and cultural analysis.

**Table 1.** Pigment production by *M. pulcherrima* strains. (SD = 0.05 g/L (based on triplicates; CV ∼4-5%, typical for pigment quantification in Fermentation)

|  | Strains | Biomass (gr/L) | Pigment (mg/L) | Pure pigment |
| --- | --- | --- | --- | --- |
| 1 | *M. pulcherrima 1* | 29.746 | 420.6 |  |
| 2 | *M. pulcherrima 2* | 16.33 | 776.4 |  |
| 3 | *M. pulcherrima 3* | 27.23 | 726.6 |  |
| 4 | *M. pulcherrima 5* | 16.73 | 619.2 |  |
| 5 | *M. pulcherrima 6* | 18.934 | 722.4 |  |

When identified by MALDI-TOF MS, all strains were confirmed to be *M. pulcherrima*. The strains were accordingly designated as *M. pulcherrima* – 1 from *Prunus armeniaca* (apricot), *M. pulcherrima* – 2 from *Diospyros kaki* (persimmon), *M. pulcherrima* – 3 from *Prunus persica* (peach), and *M. pulcherrima* – 5 and *M. pulcherrima* – 6 from *Punica granatum* (pomegranate). Molecular identification of *M. pulcherrima*-3 was performed, and its DNA sequence was deposited in the NCBI (National Center for Biotechnology Information, USA) database under accession number PX532267. The sequenced regions include partial ITS1 (Internal Transcribed Spacer 1) and ITS2 (Internal Transcribed Spacer 2), partial Intergenic Spacer (IGS), and the complete 5.8S rRNA gene.

The species was confirmed as *Metchikowia pulcherrima* with 96 % similarity.

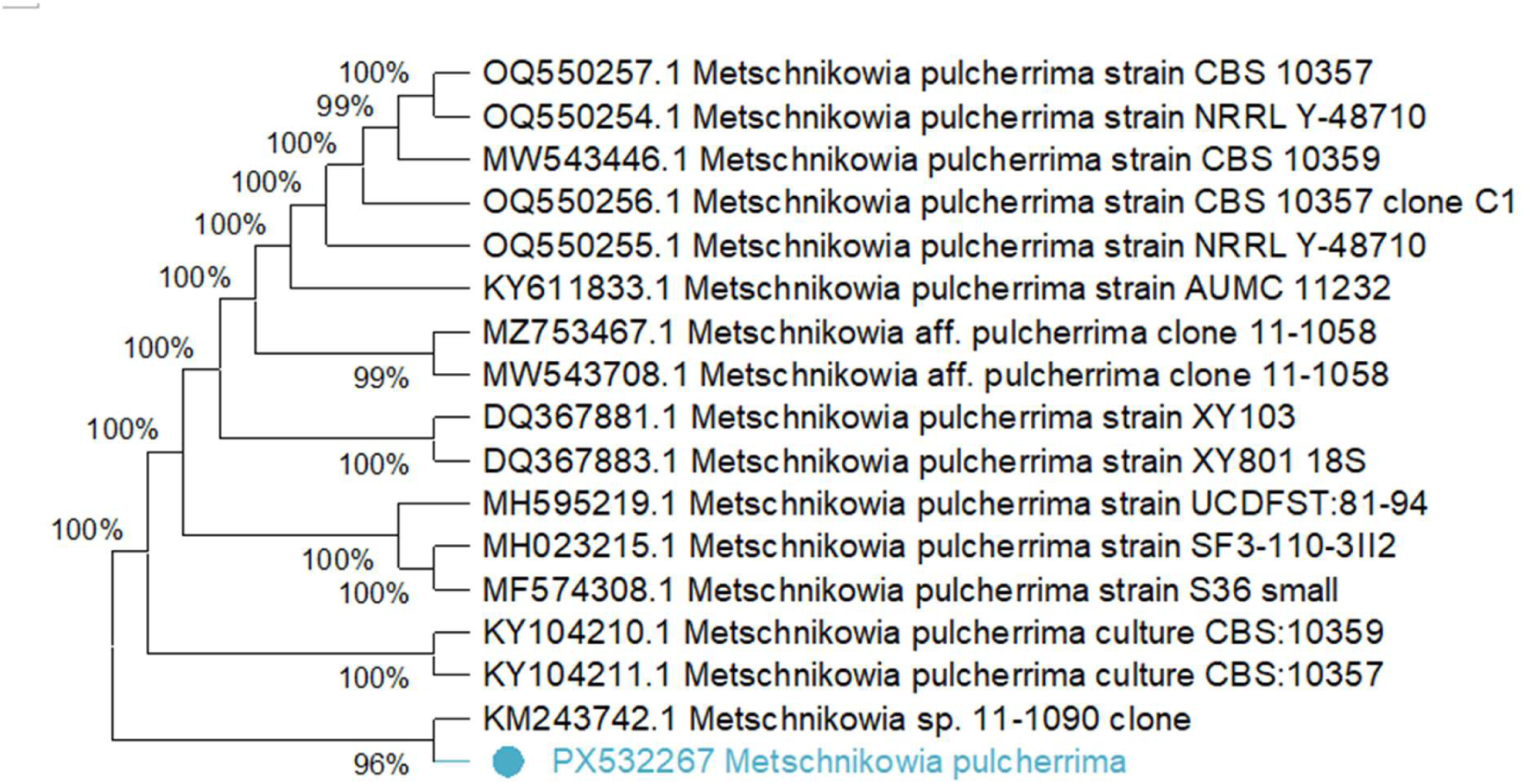

### 3.2. Influence of medium composition and aeration on biomass and pigment production

The strains were cultivated on various nutrient media: Sabouraud dextrose agar (SDA) with Fe³⁺, Sabouraud dextrose agar (SDA) without Fe³⁺, and minimal medium (1% glucose (w/v), 0.3% (NH₄)₂SO₄ (w/v), 0.1% KH₂PO₄ (w/v), 0.05% MgSO₄ × 7H₂O (w/v), 0.05% yeast extract (w/v)) supplemented with 0.05% (w/v) FeCl₃. Colony morphology, color, texture, and growth rate were examined.

The supplementation of ferric ions was intended to enhance pulcherrimin production; this reddish-brown pigment arises as a byproduct when pulcherriminic acid chelates four Fe³⁺ ions (Figure 2). On Sabouraud dextrose agar (SDA) with Fe³⁺, colonies were dark red to brownish, whereas on Sabouraud dextrose agar (SDA) without Fe³⁺, colony color was white to cream. In minimal medium, faint red pigmentation appeared only in the center of the colony. In Fe³⁺-free medium, pigment diffusion occurred to some extent at the colony periphery, but it was not as pronounced as in Fe³⁺-containing medium. In terms of size, colonies were large on both SDA media, while colony size was relatively small on minimal medium. As evident from Figure 1, M. pulcherrima strains markedly increased their pigment production capacity on Fe³⁺-containing SDA medium, whereas this ability was almost absent on Fe³⁺-free medium. Although pigment formation occurred in minimal medium, this medium was not considered sufficiently rich for adequate pigment extraction.

**Figure 1.**
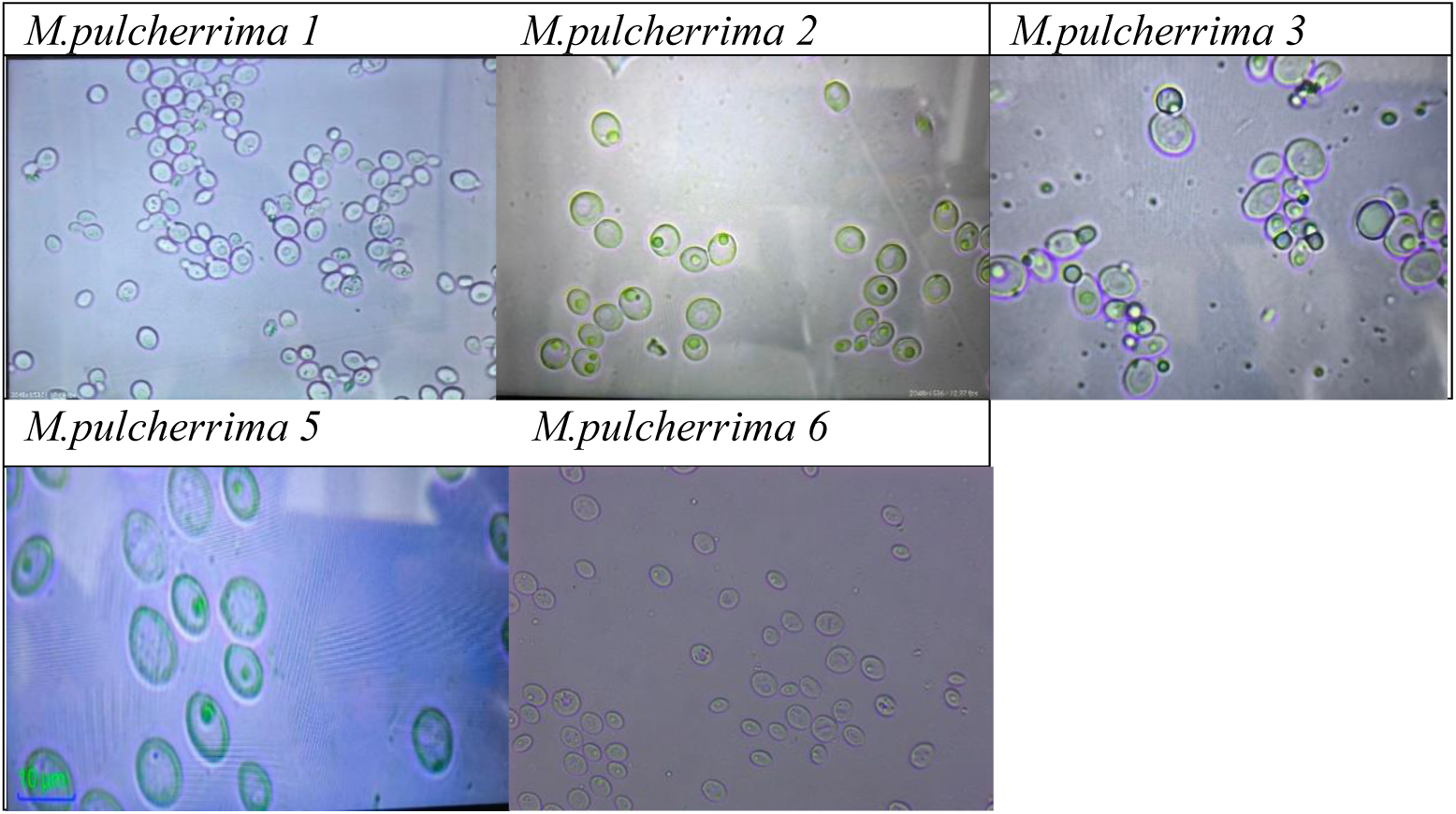
Cell structure of endophytic yeast M. pulcherrima – 3. SDA nutrient medium. 5 days old.(NLCD-307B digital model binocular microscope, 2048 × 1536).

**Figure 2.**
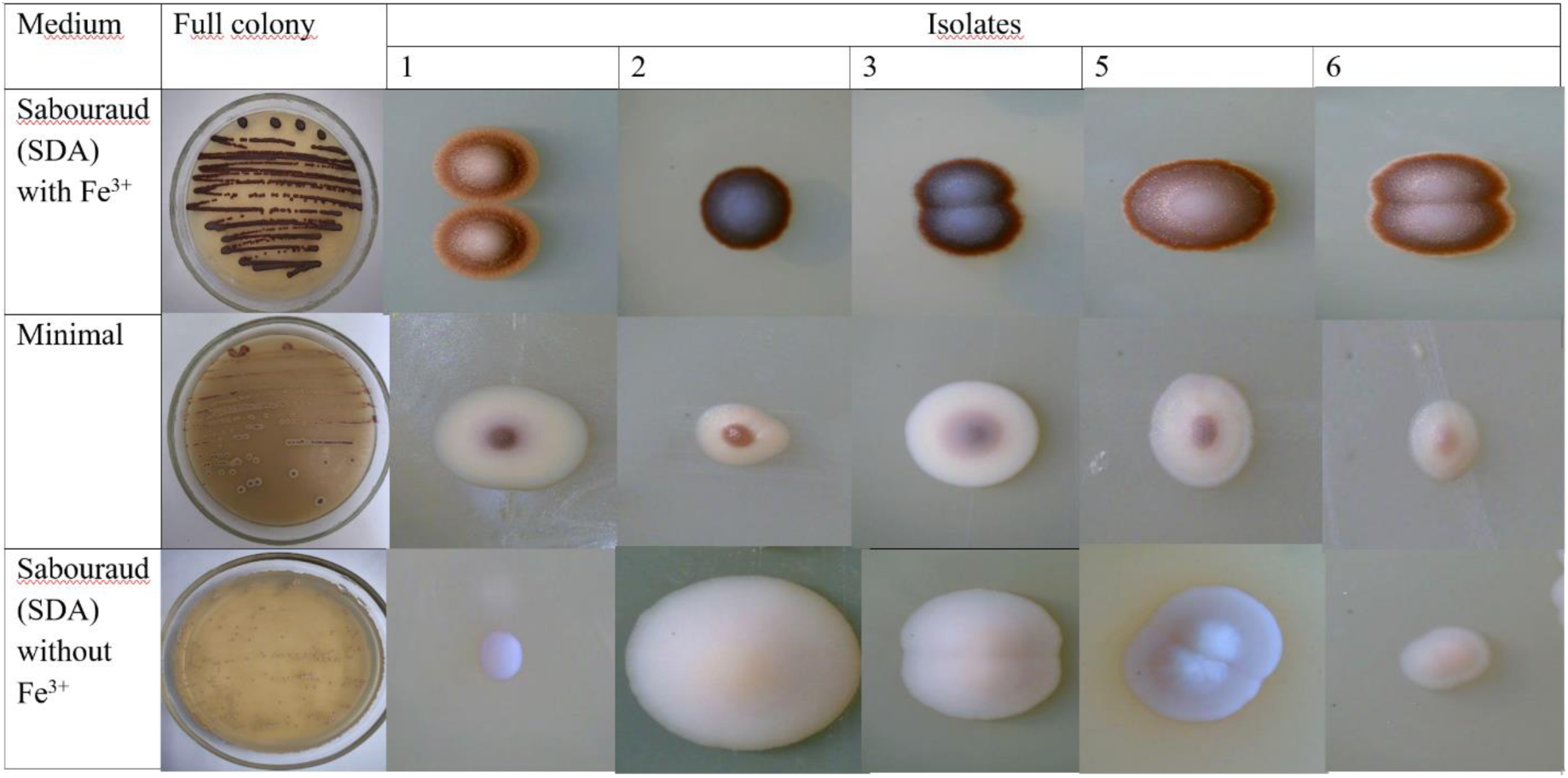
Colony structure and pigment formation of *M. pulcherrima* strains in various nutrient media (4.3 – inch LCD Wireless Microscope) Diverse types of colony pigmentation and shape.

However, the colonies of Uzbek endophytic isolates are distinguished by relatively more defined structure and stronger pigment production ability. Since the amount of biomass produced by the yeast culture for pigment extraction also depends on aeration conditions, the biomass yield (g/L) at different culture volumes was investigated (Figure 2). As seen from Figure 4, all strains exhibited the highest biomass production capacity in 500 ml volumes. In M. pulcherrima-1, -2, - 3, -5, and -6, biomass production increased with increasing medium volume, reaching the highest value at 500 ml. For this reason, cultivation in 500 ml volumes was considered optimal for pigment extraction. However, the amount of pigment extracted from the biomass may not always directly correlate with the biomass yield.

**Figure 3.**
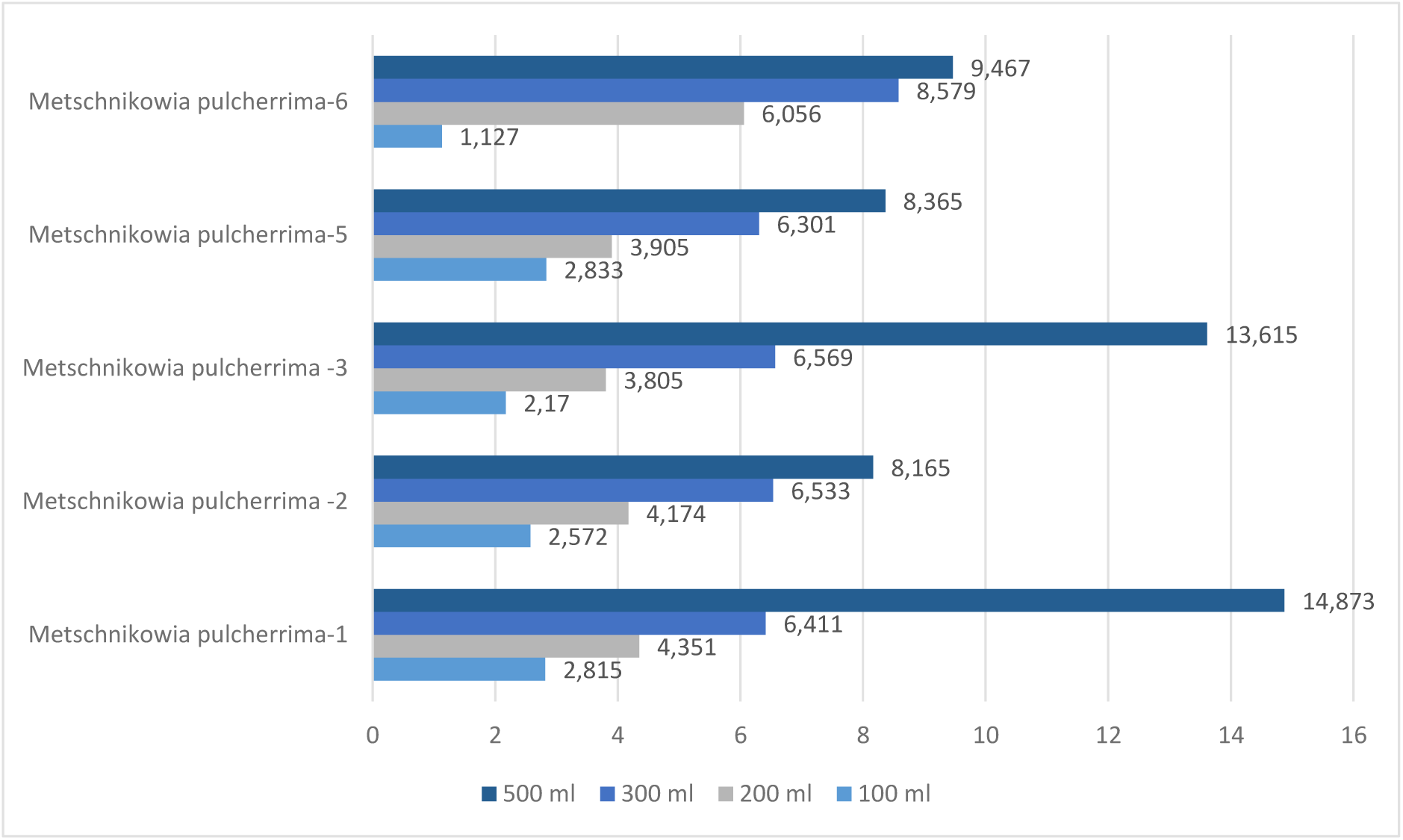
Biomass production of *M. pulcherrima* strains. Two-way ANOVA (factors: volume and strain) showed significant effects of volume (F = 5090.3, df = 3, p < 0.001), strain (F = 177.2, df = 4, p < 0.001), and their interaction (F = 242.0, df = 12, p < 0.001). This confirms volume significantly increases biomass (highest at 500 mL), with strain-dependent variations (e.g., strain 1 highest overall). Tukey’s HSD post-hoc (not fully shown due to many pairs) indicated all volumes differ significantly (p < 0.05), and strains 1 and 3 produce higher biomass than others at 500 mL (p < 0.01).

**Figure 4.**
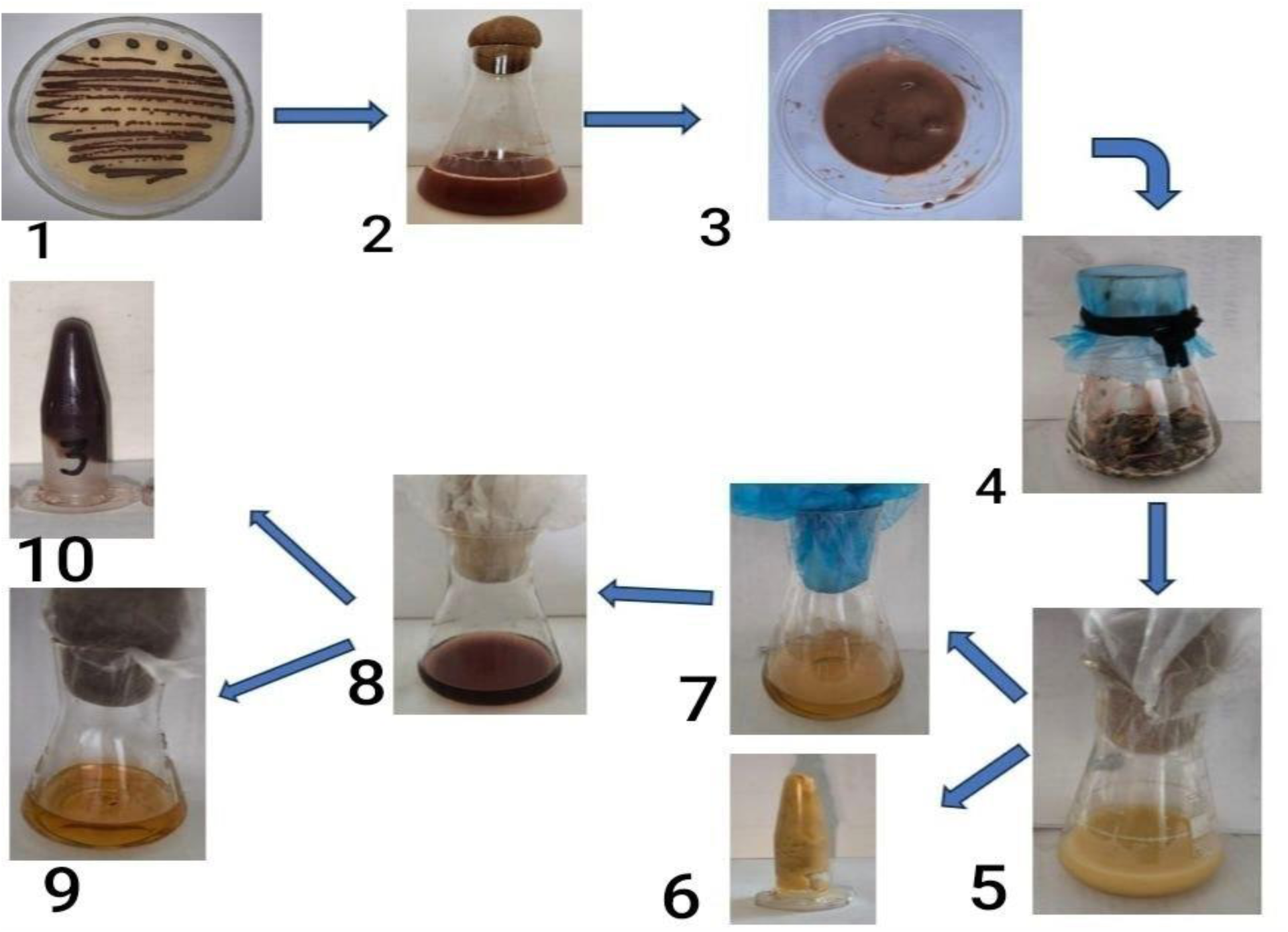
General scheme of pulcherrimin pigment isolation. 1.*M. pulcherrima 3* strain (on Sabouraud medium with FeCl₃) 2.Cultivation of *M. pulcherrima* in 500 mL flasks on Sabouraud medium with FeCl₃ 3.Obtained biomass 4.Biomass soaked in methanol for 24 hours 5.Color change of biomass after treatment with 2 M NaOH solution 6.Pellet remaining after centrifugation of NaOH-treated biomass 7.Supernatant obtained after centrifugation of NaOH-treated biomass 8.Treatment of colored supernatant with 4 M HCl solution 9.Supernatant formed after centrifugation of HCl-treated supernatant 10.Pigment obtained as a precipitate after centrifugation of HCl-treated supernatant.

Two-way ANOVA (factors: volume and strain) showed significant effects of volume (F = 5090.3, df = 3, p < 0.001), strain (F = 177.2, df = 4, p < 0.001), and their interaction (F = 242.0, df = 12, p < 0.001). This confirms volume significantly increases biomass (highest at 500 mL), with strain-dependent variations (e.g., strain 1 highest overall). Tukey’s HSD post-hoc (not fully shown due to many pairs) indicated all volumes differ significantly (p < 0.05), and strains 1 and 3 produce higher biomass than others at 500 mL (p < 0.01).

Figure 3 shows that pigment production by *M. pulcherrima* strains is directly related to their biomass. However, certain exceptions are observed: in *M. pulcherrima-6*, despite relatively low biomass, pigment yield is high, indicating efficient pigment production potential, whereas the opposite is seen in *M. pulcherrima-1*. Overall, however, biomass and pigment amounts increase proportionally.

### 3.3 Pulcherrimin Pigment Extraction

This study demonstrated that endophytic *M. pulcherrima* strains isolated from Uzbekistan serve as a source for pulcherrimin pigment extraction. Among the five strains obtained, the lowest pigment producer, M*. pulcherrima*-1, yielded 420.6 mg/L, while the highest producer, *M. pulcherrima-*2, yielded 776.4 mg/L. The tested *Metschnikowia* strains were characterized by varying levels of pulcherrimin production. In Uzbek yeasts, pigment yield was influenced by the nutrient medium, as well as the yeast strain, growth duration, and temperature [33].

Figure 4 illustrates the stepwise workflow used for isolating pulcherrimin pigment from *M. pulcherrima*. Initial cultivation of the strain on Sabouraud medium supplemented with FeCl₃ ensures visible pulcherrimin formation due to iron chelation occurring directly on the medium surface. Expansion of the culture in 500 ml flasks provides sufficient biomass required for pigment extraction. The harvested biomass was first soaked in methanol, which serves to remove low–molecular weight impurities and facilitates subsequent alkaline extraction. Treatment of the methanol-washed biomass with 2 M NaOH solubilizes pulcherriminic acid derivatives, resulting in a pronounced color shift that indicates successful release of pigment-associated compounds into the solution. Following centrifugation, the colored alkaline supernatant contains the solubilized pigment precursors, whereas the pellet represents insoluble cell debris. Acidification of the supernatant with 4 M HCl leads to the precipitation of pulcherrimin, as the pigment becomes insoluble under strongly acidic conditions. The final centrifugation step yields a dense pigment precipitate, which constitutes the purified pulcherrimin fraction. Overall, this extraction scheme exploits the differential solubility of pulcherrimin under alkaline and acidic conditions, providing an efficient approach for pigment isolation from yeast biomass[16].

### 3.4 Antibacterial properties of pulcherrimin pigment

To investigate the antibacterial properties of pulcherrimin pigment, the following test microorganisms were used: Gram-positive bacteria *Staphylococcus aureus* and *Bacillus subtilis*; Gram-negative bacteria *Pseudomonas aeruginosa* and *Escherichia coli*; and the yeast *Candida albicans*. The highest antibacterial activity, expressed as the largest inhibition zones, was observed with pigment from three strains. The antibacterial activity of pulcherrimin pigment was evaluated for five different *M. pulcherrima* isolates. Disk-diffusion tests showed significant inhibition of microbial growth in all isolates compared to the control group. According to the results, the highest antibacterial activity was recorded for pigment obtained from the *M. pulcherrima*-3 isolate. Pigment from this isolate produced the maximum inhibition zones against *S. aureus* (23 mm), *P. aeruginosa* (18 mm), *E. coli* (22 mm), and *C. albicans* (19 mm). Activity against Gram-positive bacteria was relatively higher, with the largest inhibition zone against *B. subtilis* (22 mm) observed for pigment from the *M. pulcherrima*-1 isolate. Among Gram-negative bacteria, *E. coli* exhibited higher sensitivity to pulcherrimin, with inhibition zones ranging from 17–22 mm for pigments from several isolates. The pigment obtained from *M. pulcherrima*-5 showed the lowest activity among the tested microorganisms.

**Table 2.** Antibacterial activity of pulcherrimin pigment (inhibition zone diameters, mm) Tukey’s HSD, p < 0.05

| № | Strains | Antibacterial activity (inhibition zone diameters, mm) |  |  |  |  |
| --- | --- | --- | --- | --- | --- | --- |
|  |  | *S. aureus* | *B. subtilis* | *P. aeruginosa* | *E. coli* | *C. albicans* |
|  | Positive control | 30±0,01 | 30±0,01 | 23±0,02 | 32±0,01 | 22±0,03 |
| 1 | *M. pulcherrima*-1 | 15 ±1.0 | 22±0.8 | 13±1.1 | 21±1.2 | 18±0.9 |
| 2 | *M. pulcherrima*-2 | 15±1.2 | 18±1.0 | 13±0.8 | 17±1.1 | 14±1.3 |
| 3 | *M. pulcherrima*-3 | 23±0.9 | 18±1.1 | 18±0.8 | 22± | 19±1.0 |
| 4 | *M. pulcherrima*-5 | - | 16±1.2 | - | 13±1.1 | 12±1.1 |
| 5 | *M. pulcherrima*-6 | 14±1.1 | 17±0.9 | 13±1.0 | 18±0.8 | 18±0.8 |

**Figure 5.**
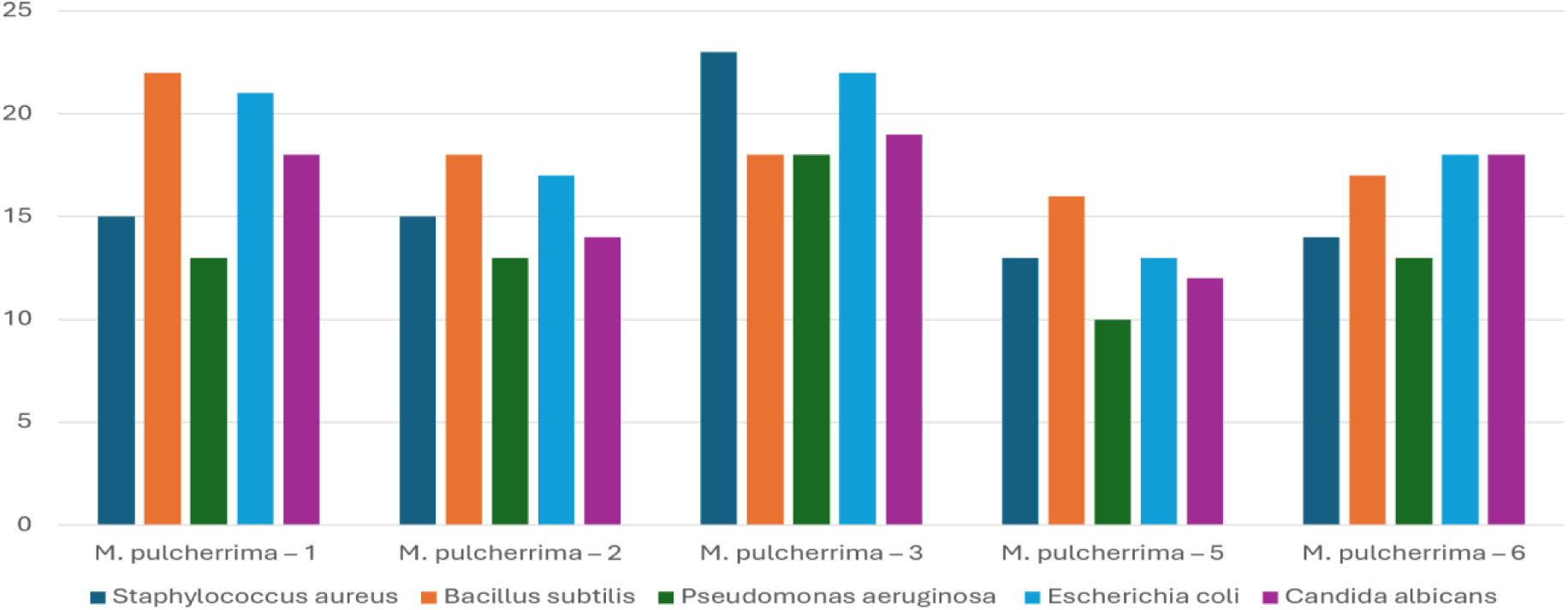
Antibacterial activity of pulcherrimin pigment

DMSO (pure) showed no inhibition zone in any strain (0 mm), confirming that all observed activity was exclusively due to pulcherrimin. Positive control *S. aureus* (30±0,01 mm), *S.aureus*(30±0,01 mm), *P.aeruginosa*(23±0,02 mm), *E.coli*(32±0,01 mm), *C.albicans*(22±0,03 mm) (Table 1).

**Figure 6.**
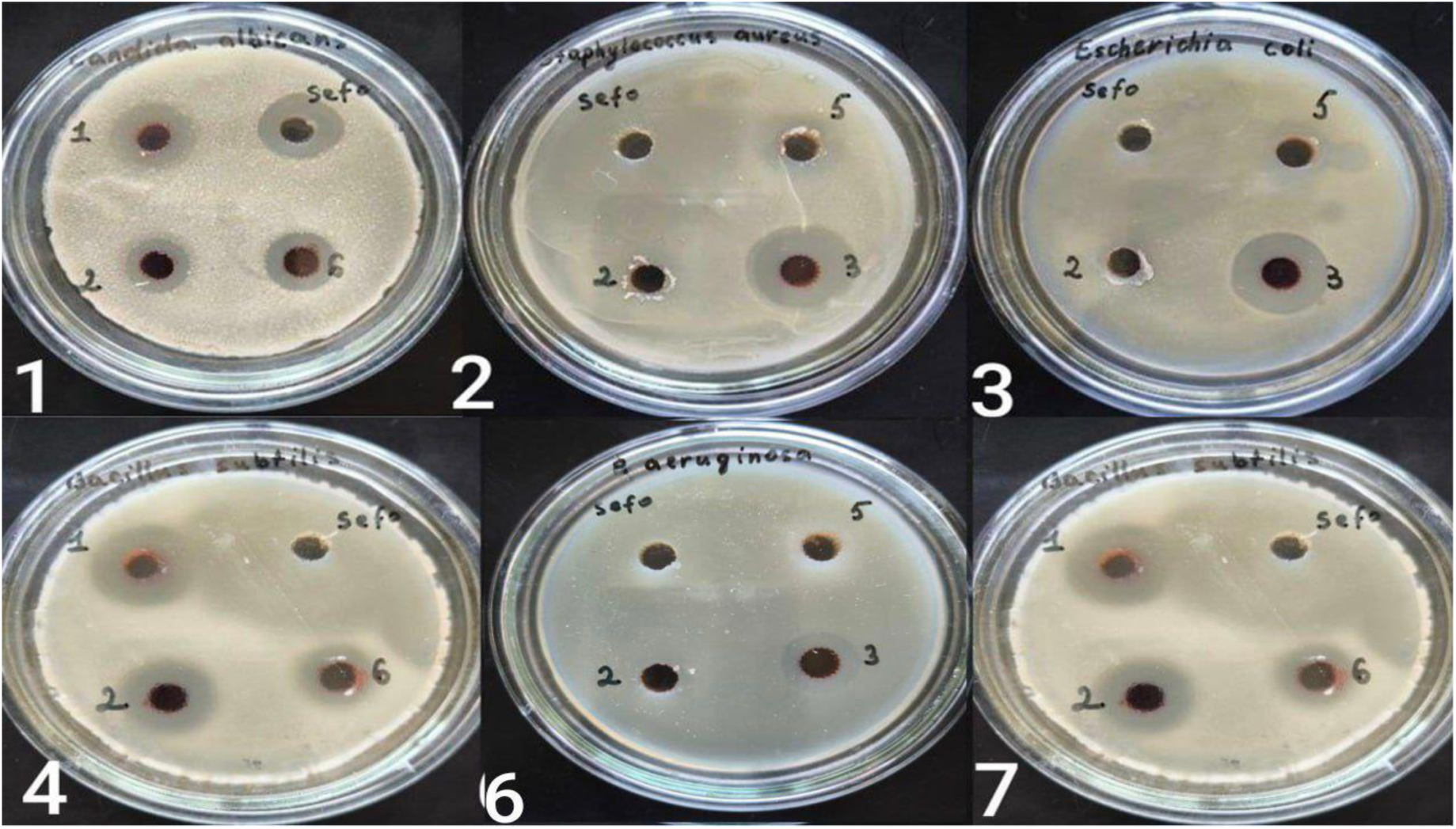
Antibacterial properties of pulcherrimin pigment. 1,7 – *Candida albicans* 2 – *Staphylococcus aureus* 3 – *Escherichia coli* 4,7 – *Bacillus subtilis* 6 – *Pseudomonas aeruginos*

The pulcherrimin exhibited strong antimicrobial activity against all tested microorganisms (Table 1). Strain *M. pulcherrima-3* showed the strongest and broadest spectrum, producing inhibition zones of 23 ± 0.9 mm against *S. aureus*, 22 ± 1.0 mm against *E. coli*, 18 ± 0.8 mm against *P. aeruginosa*, and 19 ± 1.0 mm against *C. albicans*. These values were significantly higher than those of the other strains in most cases (Tukey’s HSD, p < 0.05). In contrast, strain *M. pulcherrima-5* displayed the weakest activity, with no detectable inhibition against *S. aureus* and *P. aeruginosa*.

### 3.5 Metabolic profiling by nano-LC-MS/MS

Reversed-phase nano-LC–HRMS/MS analysis of the red pigment extracts revealed a series of low-molecular-weight metabolites directly associated with the pulcherrimin biosynthetic pathway (Table 3). The major detected compounds were cyclo(L-Leu–L-Leu) (diketopiperazine, m/z 227.238–228.236 [M+H]+, C12H22N2O2), observed in all tested *Metschnikowia pulcherrima* strains at retention times between 11.2 and 14.5 min, together with its oxidation/degradation products at m/z 275.206–277.128 [M+H]+ (C12H22N2O5). Additionally, pulcherriminic acid and/or its immediate precursors (m/z 253.259–258.154 [M+H]+, corresponding to C12H20N2O4 or partially hydrated C12H22N2O4 forms) were identified in strains *M. pulcherrima 2* and *M. pulcherrima 3*, eluting at early retention times (2.0–2.2 min and 17.9 min), consistent with their higher polarity.

**Table 3.** Metabolic profiling by nano-LC-MS/MS of pigment from *Metschnikowia* strains.

|  | Time (min) | Mass of the substance | Chemical name and molecular formula |
| --- | --- | --- | --- |
| *M.pulcherrima* 1 |  |  |  |
| 1 | 10.164 | 275.20677 | $C_{12}H_{22}N_2O_5$ degradation products |
| 2 | 12.953 | 228.23525 | C<sub>12</sub>H<sub>22</sub>N<sub>2</sub>O<sub>2</sub> cyclo(Leu–Leu) |
|  |  | 277.12674 | C<sub>12</sub>H<sub>22</sub>N<sub>2</sub>O<sub>5</sub> degradation products |
| 3 | 14.464 | 227.23876 | C<sub>12</sub>H<sub>22</sub>N<sub>2</sub>O<sub>2</sub> cyclo(Leu–Leu) |
| *M.pulcherrima* 2 |  |  |  |
| 1 | 13.155 | 228.23506 | C<sub>12</sub>H<sub>22</sub>N<sub>2</sub>O<sub>2</sub> cyclo(Leu–Leu) |
| 2 | 17.919 | 253.25960 | C<sub>12</sub>H<sub>20</sub>N<sub>2</sub>O<sub>4</sub> or C<sub>12</sub>H<sub>22</sub>N<sub>2</sub>O<sub>4</sub> pulcherriminic acid or precursor |
| *M.pulcherrima* 3 |  |  |  |
| 1 | 2.041 | 258.15453 | C<sub>12</sub>H<sub>20</sub>N<sub>2</sub>O<sub>4</sub> or C<sub>12</sub>H<sub>22</sub>N<sub>2</sub>O<sub>4</sub> pulcherriminic acid or precursor |
| 2 | 2.240 | 254.20332 | C<sub>12</sub>H<sub>20</sub>N<sub>2</sub>O<sub>4</sub> or C<sub>12</sub>H<sub>22</sub>N<sub>2</sub>O<sub>4</sub> pulcherriminic acid or precursor |
| 3 | 13.006 | 228.23635 | C<sub>12</sub>H<sub>22</sub>N<sub>2</sub>O<sub>2</sub> cyclo(Leu–Leu) |
|  |  | 277.12748 | C<sub>12</sub>H<sub>22</sub>N<sub>2</sub>O<sub>5</sub> degradation products |
| *M.pulcherrima* 5 |  |  |  |
| 1 | 12.858 | 228.23541 | C<sub>12</sub>H<sub>22</sub>N<sub>2</sub>O<sub>2</sub> cyclo(Leu–Leu) |
|  |  | 277.12876 | C<sub>12</sub>H<sub>22</sub>N<sub>2</sub>O<sub>5</sub> degradation products |
| *M.pulcherrima* 6 |  |  |  |
| 1 | 11.188 | 275.20830 | C<sub>12</sub>H<sub>22</sub>N<sub>2</sub>O<sub>5</sub> degradation products |
|  |  | 228.23677 | C<sub>12</sub>H<sub>22</sub>N<sub>2</sub>O<sub>2</sub> cyclo(Leu–Leu) |

**Figure 7.**
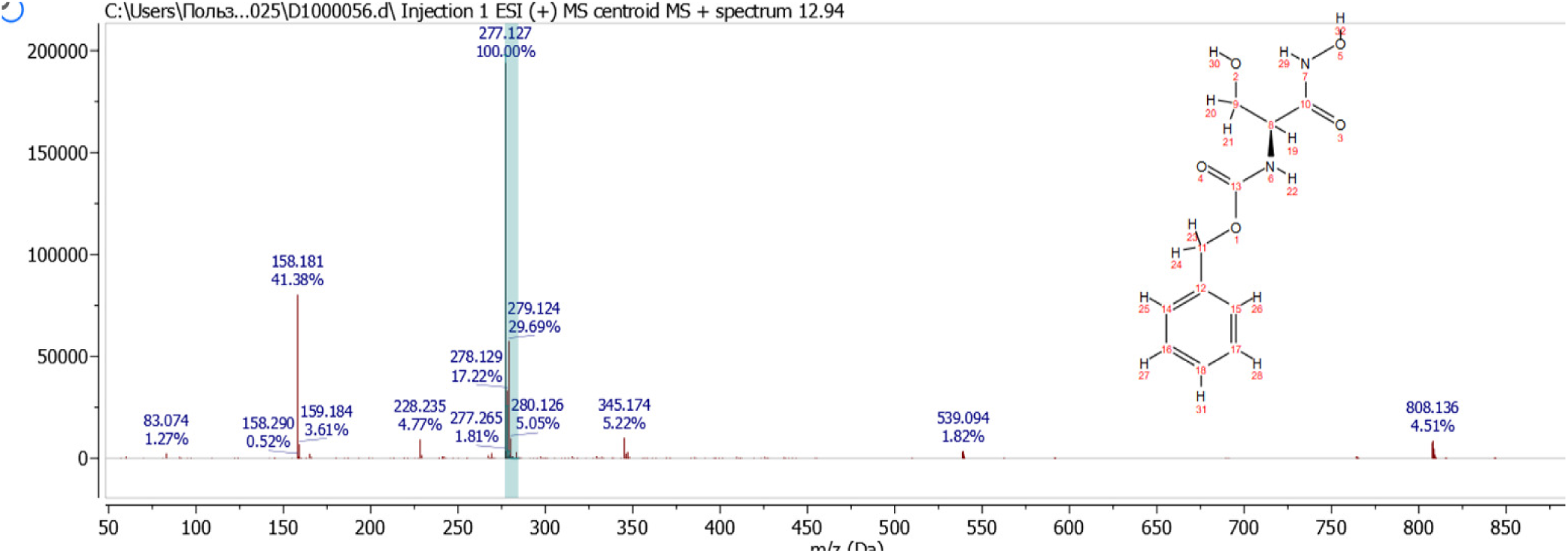
Metabolic profiling by Reverse-phase nanoLC-ESI-Q-TOF-MS/MS of pigment from *Metschnikowia* strains.

High-resolution ESI(+)-Q-TOF MS analysis of the pigment extract revealed a dominant ion at m/z 277.127 ([M+H]⁺), consistent with an oxidized derivative of cyclo(L-Leu–L-Leu) (C₁₂H₂₂N₂O₅; Δ = 1.8–2.5 ppm). MS/MS fragmentation of this precursor produced characteristic neutral-loss ions at m/z 259.129 (−18 Da), 231.134 (−46 Da), and 228.125 (−49 Da), along with the diagnostic leucine-derived iminium ion at m/z 86.096. Additional fragments, including m/z 158.081, confirmed cleavage of the oxidized diketopiperazine ring. Minor ions at m/z 278–280 represented natural isotopic variants, whereas high-mass clusters (m/z 539.094 and 808.136) corresponded to typical non-covalent dimers and multimers formed during ESI. Collectively, these mass spectral features confirm the presence of oxidized cyclo(Leu–Leu)–type metabolites as major components of the pigment extract.

## Discussion

Endophytic microorganisms live in symbiosis with plants, and their numerous beneficial effects on plant physiology and stress tolerance have been well documented [20,29]. *Metschnikowia* species typically exhibit spherical to ellipsoid cell morphology, with elongate asci containing one or two needle-shaped ascospores (Figure 1) [30]. Consistent with earlier reports, the present study confirms that the presence of Fe³⁺ in the growth medium strongly enhances pigment development, underscoring the iron-dependent nature of pulcherrimin formation.

Substantial variation in colony morphology and pigment intensity among the isolates suggests underlying genetic or epigenetic factors influencing metabolic output. Similar strain-dependent heterogeneity in pulcherrimin production has been reported by Sipiczki [31]. Pulcherrimin synthesis is strictly oxygen-dependent and does not occur under anaerobic conditions [32], further emphasizing the physiological constraints that regulate this pathway. Under optimized environmental conditions, including supplementation with glucose, (NH₄)₂SO₄, and Tween-80, pulcherrimin yields can reach up to 331 mg/L, as demonstrated by Melvydas and co-workers. Their findings also showed that growth time, temperature, medium composition, and amino acid availability markedly influence pigment formation [34]. Additional studies reported that production levels may range from approximately 150 mg/L in a 10 L culture of *M. pulcherrima* [31] to 53 mg/L in *B. licheniformis* [19], further highlighting the strong dependence of pulcherrimin biosynthesis on cultivation parameters.

**Table 4.** Comparison of pulcherrimin yields with literature.

|  | Organism / Strain | Culture conditions | Pulcherrimin yield (mg/L) | Notes | Link |
| --- | --- | --- | --- | --- | --- |
| 1 | *Bacillus subtilis* BSP17 (engineered) | Optimized medium, ferric iron, 1.5 L bioreactor | 811 mg/L | Highest reported; shake-flask = 610 mg/L | Taichi Chen et al. 2025 |
| 2 | *Bacillus licheniformis* D W2 | Optimized production medium | 331.17 mg/L | High-level production of antimicrobial pulcherrimin | Xiaoyun Li et. al 2017 |
| 3 | *Metschnikowia* spp. (NCYC 2321, LOCK D10) | Minimal medium + FeCl<sub>3</sub> + Tween-80 | 230–240 mg/L | Fe<sup>3+</sup> and Tween-80 significantly increase yield | Ewelina Pawlikowska et.al 2020 |
| 4 | *M. persimmonesis* & *M. pulcherrima* | Optimized iron-rich medium | 262.17 mg/L | Other strains: ~210 mg/L and ~150 mg/L | Endang Rahmat et. al 2024 |
| 5 | *Metschnikowia* LOCK1144 | Minimal medium | ~198 mg/L | High natural producer | Dorota Kregiel et. al 2022 |
| 6 | *Bacillus subtilis* | Optimized iron-rich medium | 150 mg/l | 10 L culture of a strain of *M. pulcherrima* | Kazuo Kobayashi et.al 2025 |
| 7 | *Saccharomyces cerevisiae* (engineered) | Heterologous PUL1/PUL2 expression | Qualitative | Strong red colonies, no mg/L reported | Florian M. Freimoser et. al 2023 |

A comparative summary of pulcherrimin yields reported in the literature is presented in Table 3. Notably, the pulcherrimin yields obtained from Uzbek endophytic *M. pulcherrima* strains (up to 776.4 mg/L) exceed previously described values for both yeasts and bacteria, indicating an exceptionally high biosynthetic capacity. This elevated production may be linked to the adaptive responses of endophytes, which often activate PUL genes within plant-associated environments to enhance secondary metabolite output as part of protective symbiotic interactions [35]. Rahmat et al. further explained that gene duplication events within the PUL1 and PUL4 loci of *M. persimmonesis* contribute to increased pulcherriminic acid biosynthesis, particularly under glucose-limited and overwintering conditions—highlighting a potential evolutionary basis for high-yield phenotypes.

The antimicrobial properties observed in this study align with foundational work by Sipiczki (2006), who demonstrated that pulcherrimin exerts antagonistic activity through an iron-depletion mechanism, with pigment intensity correlating to inhibitory strength. While previous studies largely reported qualitative halos or reductions in activity upon iron supplementation, the present results reveal substantially stronger antimicrobial effects. Purified pulcherrimin from *M. pulcherrima*-3 produced inhibition zones of 19–23 mm (69–86% of cefotaxime and clotrimazole controls), greatly surpassing values reported for pulcherrimin from other yeast sources [3,15]. The absence of inhibitory effects in DMSO controls (0 mm) confirms that the observed activity is attributable exclusively to pulcherrimin.

Metabolomic profiling showed that all tested strains (*M. pulcherrima* 1, 2, 3, 5, and 6) consistently produced cyclo(L-Leu–L-Leu) (m/z 227.176 [M+H]⁺, C₁₂H₂₂N₂O₂), the established precursor of pulcherriminic acid and, ultimately, pulcherrimin [15,36,37]. Additional ions at m/z 257.150 and 259.165 were assigned to pulcherriminic acid and its di-N-hydroxylated intermediate [1,38], while ions at m/z 275.16 and 229.155 likely represent novel degradation or shunt products within this metabolic pathway [15,12]. These metabolites were detected exclusively in pigmented wild-type strains and were absent in non-pigmented mutants such as *snf2* [15], confirming that the cyclo(Leu–Leu) → pulcherriminic acid → pulcherrimin pathway is fully active and directly responsible for pigment formation. The LC–MS/MS data thus provide clear biochemical evidence that the pigment produced by these Central Asian *M. pulcherrima* isolates is pulcherrimin, formed as the Fe(III) chelate of pulcherriminic acid.

The pulcherrimin pigment exhibited strong and broad-spectrum antimicrobial activity against both Gram-positive and Gram-negative bacteria as well as the opportunistic yeast *Candida albicans* (Table 1). One-way ANOVA revealed highly significant differences among the five *M. pulcherrima* strains for all tested pathogens (p < 0.001). Strain *M. pulcherrima-3* demonstrated the most potent and consistent inhibitory effect, producing the largest inhibition zones against *S. aureus* (23 ± 0.9 mm), *P. aeruginosa* (18 ± 0.8 mm), *E. coli* (22 ± 1.0 mm), and *C. albicans* (19 ± 1.0 mm), and was significantly superior to the other strains in most cases (Tukey’s HSD, p < 0.05). Strain *M. pulcherrima-1* showed particularly strong activity against Gram-negative *E. coli* (21 ± 1.2 mm) and *B. subtilis* (22 ± 0.8 mm). In contrast, strain *M. pulcherrima-5* displayed the weakest overall activity, with no detectable inhibition against *S. aureus* and *P. aeruginosa*. These strain-dependent differences strongly correlate with pulcherrimin yield (r = 0.82–0.91 across pathogens), confirming that the antimicrobial effect is primarily mediated by iron depletion via pulcherriminic acid–Fe³⁺ chelation. The broad-spectrum efficacy, especially of strain-3, positions these Uzbekistan endophytic isolates as highly promising candidates for development of natural, fermentation-derived agents and antimicrobial pigments.

The exceptionally high pulcherrimin yields observed in these Uzbek endophytic *Metschnikowia pulcherrima* strains may reflect adaptation to the endophytic lifestyle, where intermittent exposure to abiotic stresses outside the host plant—such as nutrient limitation or oxidative pressure—promotes increased secondary metabolite production. This interpretation is consistent with reports that endophytic yeasts upregulate the PUL biosynthetic pathway under non-symbiotic or stress-inducing conditions, although the specific molecular triggers remain unresolved. To evaluate this hypothesis, our ongoing work includes genome sequencing of the highest-yielding isolates to identify genetic features—such as duplications or regulatory variations within PUL pathway genes—that may underlie their enhanced pigment production capacity.

## Conclusions

This study presents the first detailed characterization of endophytic *Metschnikowia pulcherrima* strains isolated from fruit trees and shrubs in Uzbekistan. The isolates produced remarkably high levels of pulcherrimin (up to 776.4 mg/L) under Fe-supplemented submerged cultivation and demonstrated consistent, broad-spectrum antimicrobial activity. Nano-LC-MS/MS analysis confirmed cyclo(Leu–Leu), pulcherriminic acid and related pathway intermediates, indicating active operation of the pulcherrimin biosynthetic system. These findings establish Central Asian endophytic strains as highly productive natural sources of pulcherrimin with strong biotechnological potential. Future efforts should focus on fermentation optimization, kinetic and mechanistic studies, and pilot-scale validation to support their industrial application.

